# A Spatiotemporal History of Key San Francisco Estuary Pelagic Fish Species

**DOI:** 10.1101/2022.08.26.505491

**Authors:** Dylan K. Stompe, Peter B. Moyle, Kiva L. Oken, James A. Hobbs, John R. Durand

## Abstract

Estuaries across the globe have been subject to extensive abiotic and biotic changes and are often monitored to track trends in species abundance. The San Francisco Estuary is a novel ecosystem that has been deeply altered by anthropogenic factors, resulting in fish declines over the past 100 years. To track these species declines, a patchwork of monitoring programs has operated regular fish surveys dating back to the late 1950s. While most of these surveys are designed to track population-scale changes in fish abundance, they are methodologically distinct, with different target species, varying spatial coverage and sample frequency, and differing gear types. To remediate for individual survey limitations, we modeled pelagic fish distributions with integrated data from many sampling programs. We fit binomial generalized linear mixed models with spatial and spatiotemporal random effects to map annual trends in the distribution of detection probabilities of striped bass, Delta smelt, longfin smelt, threadfin shad, and American shad for the years 1980 to 2017. Detection probabilities decreased dramatically for these fishes in the Central and South Delta, especially after the year 2000. In contrast, Suisun Marsh, one of the largest tidal marshes on the west coast of the United States, acted as a refuge habitat with reduced levels of decline or even increased detection probabilities for some species. Our modeling approach demonstrates the power of utilizing disparate datasets to identify regional trends in the distribution of estuarine fishes.

## Introduction

Estuaries are highly productive and often urbanized systems located at the interface between freshwater and marine environments. Biological productivity is fueled through the input of both terrestrial and marine nutrients, as well as through increased residence time due to salinity-driven density gradients and tidal forcing (Pérez-Ruzafa et al. 2011). Estuaries support diverse assemblages of aquatic species along the salinity gradient, from obligate stenohaline species at the marine and freshwater fringes to oligohaline species that can utilize the entirety of the estuary (Whitfield et al. 2022). The high productivity and location of estuaries at the interface between freshwater and marine environments has also resulted in the general colonization of estuaries by humans seeking the benefit of rich food sources and protected ports (Wilson 1988; Lotze et al. 2006; Cabral et al. 2022).

The dynamic nature of estuaries has encouraged, and in some cases necessitated, environmental modification for human habitation. Many large estuaries have been diked, drained, and dredged for urbanization, transport, water management, flood control and agriculture. Inflows have been diverted or impounded in reservoirs (Cabral et al. 2022). These changes have substantially altered abiotic conditions and have in some cases increased the habitability of estuaries to non-native species (Cabral et al. 2022; Moyle and Stompe 2022.). Species introductions are common in estuaries through ballast water exchange of hitchhiking organisms from international shipping traffic. Recreational, ornamental, and bait species are also often introduced (Moyle and Stompe 2022).

Estuaries are frequently monitored to track trends in the abundance and distribution of aquatic species of recreational, commercial, and cultural importance, many of which have declined because of extensive abiotic and biotic changes (Blaber et al. 2022; Cowley et al. 2022). Estuarine monitoring is often undertaken by state, federal, tribal, academic, and/or non-governmental organizations, all with potentially different objectives (Anderson 2005). The often-piecemeal implementation of surveys complicates traditional analyses of species trends. In some cases, this results in underutilization of data due to concerns about differences in methodology and bias (Stompe et al. 2020; Huntsman et al. 2022). Unfortunately, logistical and/or financial limitations of surveys means that trends in abundance and distribution of estuarine species are often incomplete when based on analysis of a single survey.

The San Francisco Estuary (Estuary) is a well-monitored system in which data resources have not been fully utilized. The Estuary includes the tidally influenced portions of the Sacramento and San Joaquin Rivers, the Delta, Suisun Bay and Marsh, San Pablo Bay, Central and Southern San Francisco Bay; it terminates at the Golden Gate (Figure 1). Habitats within the Estuary are diverse and include pelagic marine habitats, salt, brackish, and freshwater marshes, low gradient riverine habitats, and freshwater littoral habitats, among others.

**Figure 1.**
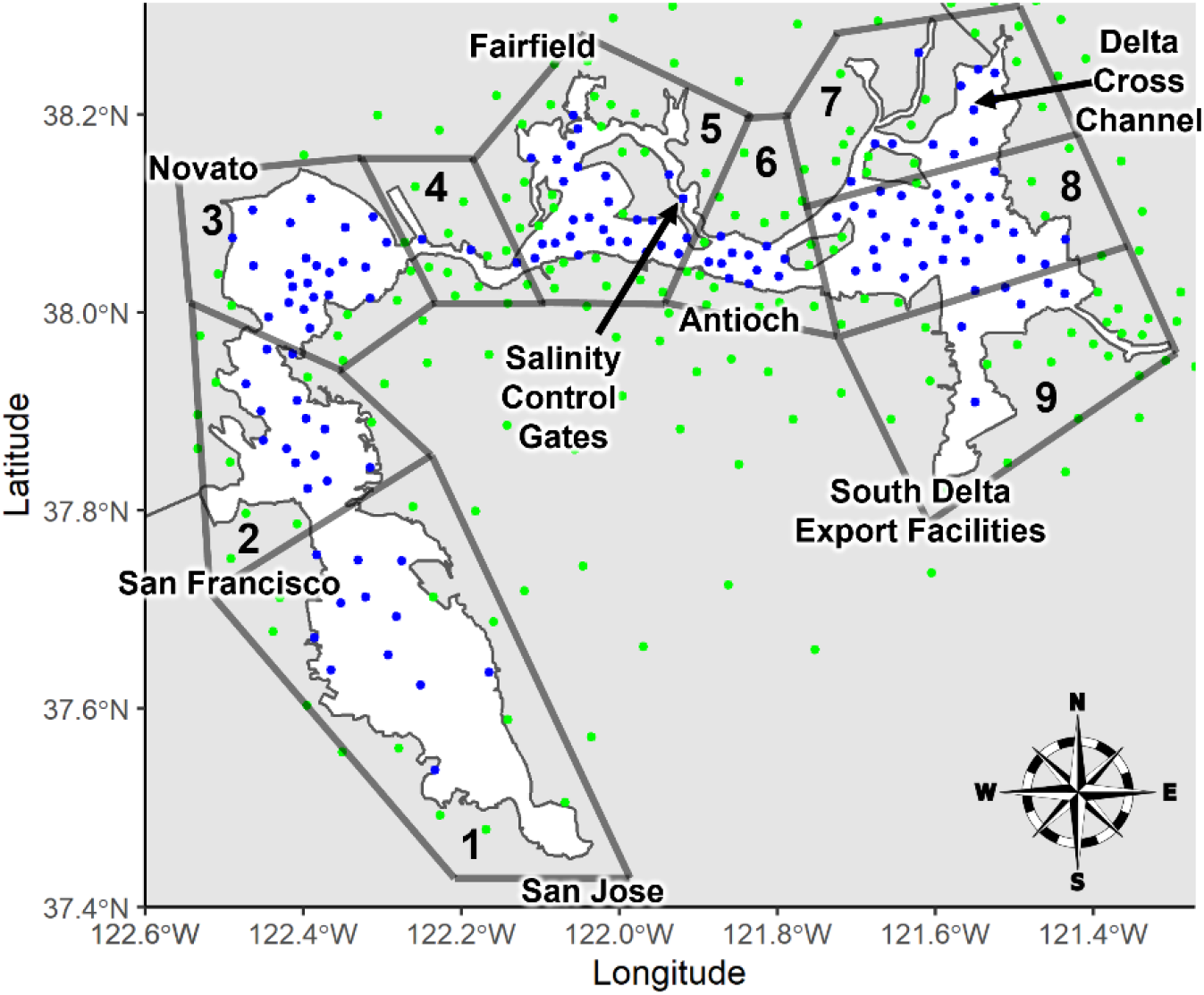
Simplified spatial plane of the San Francisco Estuary with applied barrier components. White background is wetted area and grey background is land. Blue dots represent the center of “water” spatial mesh triangles and green dots represent the center of “land” spatial mesh triangles. Select cities surrounding the estuary, the location of the Montezuma Slough salinity control gates, the location of the Delta Cross Channel, and the location of the South Delta export facilities (State Water Project, Central Valley Project) are included for reference. Numbered regions are identified for regional descriptions of distribution trends. 1 = South San Francisco Bay, 2 = Central San Francisco Bay, 3 = San Pablo Bay, 4 = Carquinez Strait, 5 = Suisun Marsh (top) and Suisun Bay (bottom), 6 = Sacramento-San Joaquin Confluence, 7 = North Delta, 8 = Central Delta, 9 = South Delta.

The Estuary has been highly altered by myriad abiotic and biotic changes since large-scale colonization by European-Americans in the 1800s. Abiotic changes include extensive physical and hydrologic alterations in the form of water diversions, levees, floodplain and marshland reclamation, construction of dams on every major river in the Estuary’s 163,000 km^2^ watershed, and the severe disruption of natural sedimentation regimes due to historic hydraulic mining and the current impoundment of sediment behind dams (Cloern and Jassby 2012; Whipple et al. 2012; Schoellhamer et al. 2013; Herbold et al. 2014). Sometimes called the most heavily invaded estuary in the world, the Estuary has also been subject to numerous species invasions through ballast water dumping in its freshwater and saltwater ports, and state-sponsored and illicit intentional introductions (Cohen and Carlton 1998). As a result of these changes, and other stressors such as overfishing (Yoshiyama et al. 1998) and pollution (Brooks et al. 2012), many native and some introduced fish species have experienced drastic declines in the past 100 years, and especially the last 40 years (Sommer et al. 2007; Stompe et al. 2020).

Along with numerous smaller diversions, the Estuary contains two major water export facilities located in the South Delta, the State Water Project (SWP) and Central Valley Project (CVP; Figure 1), which export a large proportion of freshwater inflow for agricultural and municipal use (Gartrell et al. 2017; Moyle et al. 2018). To track the impacts of these water infrastructure operations on Estuary fish species, state and federal agencies and a research group at the University of California, Davis (UC Davis), have operated regular fish surveys dating back to the late 1950s (Honey et al. 2004; Herrgesell 2012; Tempel et al. 2021). Early surveys primarily focused on tracking the abundance of young of year striped bass, an introduced yet recreationally and culturally important species within the Estuary (Turner and Chadwick 1972; Chadwick et al. 1977; Stevens et al. 1985). Later, additional surveys were started to track the abundance and survival of outmigrating juvenile salmonids (Dekar et al. 2013) and to track changes in fish and invertebrate assemblages (Baxter et al. 1999; O’Rear et al. 2021) rather than single species.

While most Estuary fish surveys are designed to track fish abundance over a wide spatial expanse, they are logistically and economically restricted in total number of stations and frequency of sampling events. In addition, surveys sample different micro-and macro-habitats due to differences in gear type, spatial coverage, and project goals (Tempel et al. 2021). Since fish species are not homogenous in their distribution throughout the water column or Estuary, differences in catch arise because of these disparities (Stompe et al. 2020).

In this paper, we leverage an integrated dataset of eight long-term Estuary surveys (Stompe et al. 2020) to examine trends in the distribution and abundance of several important fish species. Species included are striped bass (*Morone saxatilis*), Delta smelt (*Hypomesus transpacificus*), longfin smelt (*Spirinchus thaleichthys*), threadfin shad (*Dorosoma petenense*), and American shad (*Alosa sapidissima*). We use spatially-explicit species distribution modeling to fit binomial generalized linear mixed models to the integrated survey data and then 1) identify key areas of importance for these species over time, 2) pinpoint time periods of major shifts in distribution and abundance, and 3) describe the effects of freshwater outflow on distribution.

## Methods

### Study Species

The species selected for our analysis (striped bass, Delta smelt, longfin smelt, threadfin shad, American shad) were chosen due to their ecological, cultural, and recreational significance in the Estuary. These species represent important recreational fisheries (striped bass and American shad) and both native (Delta smelt, longfin smelt) and introduced (threadfin shad) forage fishes. In addition, all these fishes require productive estuarine pelagic environments during part or all their lives (Moyle 2002), so their abundances can be indicators of the ‘health’ of pelagic habitats.

Striped bass is an introduced, relatively long-lived, semi-anadromous, and fecund species that relies on productive estuaries for rearing (Raney 1952). Since their introduction, they have been one of the primary catch species in agency and university surveys, although catches have declined considerably over the years (Kohlhorst 1999; Sommer et al. 2007).

Delta smelt is a small native osmerid endemic to the Estuary. They are generally an annual species and are an obligate estuarine species (Moyle et al. 1992; Moyle 2002). Despite once high abundance, they are now rarely caught by Estuary surveys and are listed as threatened under the Federal Endangered Species Act (FESA) and endangered under the California Endangered Species Act (CESA; Tempel et al. 2021).

Like Delta smelt, longfin smelt are small native osmerids; they are found along the Pacific Coast of North America, are more halophilic than Delta smelt, and can live two to three years (Moyle 2002). Historically, they were highly abundant within the Estuary but have since declined and are now relatively rare (Sommer et al. 2007). As a result, they were listed as threatened under the CESA in 2009 (Tempel et al. 2021).

Threadfin shad are introduced, small, deep bodied clupeids that typically live two to three years (Moyle 2002). Despite their somewhat recent introduction to the system, threadfin shad have also experienced declines in abundance (Sommer et al. 2007).

American shad are another introduced clupeid species, but they reach larger sizes than threadfin shad and generally migrate into the Pacific Ocean after rearing in the Estuary (Moyle 2002). Unlike the above-described fishes, previous work has not identified major reductions in Estuary American shad abundance and instead has seen recent increases in angler catch (Ferguson 2016).

### Survey Data

The surveys included in our modeling effort are the California Department of Fish and Wildlife (CDFW) Fall Midwater Trawl (FMWT; White 2021), CDFW Bay Study Otter and Midwater Trawls (BSOT, BSMT; Baxter et al. 1999), CDFW Summer Townet Survey (STN; Malinich 2020), UC Davis Suisun Marsh Otter Trawl and Beach Seine Surveys (SMOT, SMBS; O’Rear et al. 2021), United States Fish and Wildlife Service (USFWS) Beach Seine Survey (BSS; McKenzie 2021a), and the USFWS Chipps Island Trawl (CIT; McKenzie 2021b; Table 1). Of the stations we include in our analysis, there is considerable spatial overlap between the surveys in the San Pablo, Carquinez, Suisun, and Sacramento-San Joaquin Confluence Regions (Figure 1 – Regions 3-6; Appendix Figure 1). Conversely, the Bay Study Otter Trawl and Bay Study Midwater Trawl are the only surveys with stations in the Central and South San Francisco Bays (Figure 1 – Regions 1 and 2; Appendix Figure 1) and only the Fall Midwater Trawl, Summer Townet Survey, Beach Seine Survey, and Chipps Island Trawl have stations in the Delta (Figure 1 – Regions 7-9; Appendix Figure 1). The longest running of these surveys (Summer Townet Survey) started in 1959 and the most recent (Bay Study Otter and Midwater Trawls) in 1980, and all have operated continuously on at least an annual basis through 2017. Several of these surveys were originally designed to describe and track trends in young of year striped bass abundance (Fall Midwater Trawl, Summer Townet Survey), some were designed to track the outmigration of juvenile salmonids through the Estuary (Chipps Island Trawl, Beach Seine Survey), and others were designed to track assemblages of fish and invertebrate populations (Suisun Marsh Otter Trawl/Beach Seine, Bay Study Otter and Midwater Trawls). All were implemented to determine the effects of water diversion and/or entrainment into export facilities on fish populations. These surveys primarily capture small and/or juvenile fishes due to their specific gear types and netting mesh sizes. For this reason, surveys are generally able to describe trends in the relative abundance of small fishes such as Delta smelt, Longfin smelt, and threadfin shad, but only represent juvenile trends for large fishes, such as striped bass and American shad.

**Table 1.**
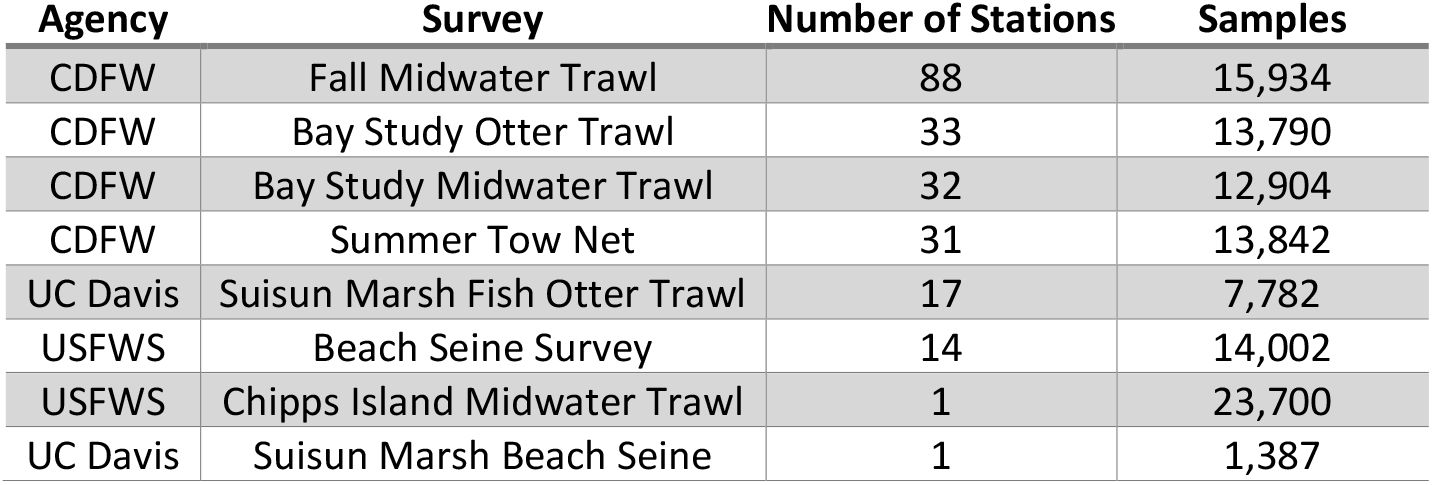
Surveys, number of stations, and total number of samples included in eight-survey dataset. Samples are indexed as individual trawl or seine pulls. CDFW = California Department of Fish and Wildlife and USFWS = United States Fish and Wildlife Service.

Data from these surveys were integrated into an aggregate dataset for the years 1980 through 2017, retaining key variables such as date, coordinates, and number of individual fish captured by species (Stompe et al. 2020). For consistency of annual spatial extent, we only include those survey stations which were sampled at least once annually in our analyses. In addition, several Beach Seine Survey stations located on the Sacramento River upstream of the Delta were omitted because of their distance from other downstream stations. Because effort shifted among certain years, data used from the Chipps Island Trawl was seasonally restricted to April through June and from the Fall Midwater Trawl to September through December to standardize seasonal effort. All other surveys generally operated year-round.

Catch per unit effort was calculated as total number of fish caught per seine or trawl, with a total of 103,341 sampling events. While other Estuary data integration efforts have instead chosen to index catch by the volume of water sampled (Huntsman et al. 2022), not all surveys we include record this metric, nor is it as meaningful of a metric for some gear types (i.e. beach seines). In addition to indexing catch per trawl or seine, we included species presence or absence for each sample. The inclusion of a binary metric allows for the modeling of probability of detection, regardless of water volume sampled.

In this integrated format, these data represent approximately 40 years of trends in Estuary fish abundance at a much greater seasonal and spatial density than could be provided by any single survey. In addition, the breadth of gear types included in the eight-survey dataset mean that benthic, pelagic, and littoral species are all targeted to some degree.

### Data Analysis

Using the aggregate eight-survey dataset, we modeled spatiotemporal trends in the probability of detection of the previously described fish species using the package ‘sdmTMB’ (Anderson et al. 2022). We constructed and fit binomial generalized linear mixed models (GLMMs) of species presence by maximum likelihood for each of the five fish species from the aggregate dataset across a restricted spatial mesh. Model predictions from the binomial models represent the probability of detection rather than the total predicted catch of a given species. Because of this, these models do not necessarily capture absolute changes in abundance. However, binomial model predictions do provide some index of changes in abundance because survey gear is more likely to detect species at higher densities assuming they are relatively evenly distributed within the sampled habitats and assuming limited density dependence of catch (Godø et al. 1999). During model construction Tweedie distributions (Tweedie 1984) were also tested, but model fit was poor with this distributional family (Hartig 2022).

A spatial mesh for efficiently modeling spatial and spatiotemporal autocorrelation was generated using the “cutoff” method, with a minimum of 2km spacing between mesh vertices (“knots”) and resulting in a mesh with 179 knots (Anderson et al. 2019). Knot spacing was iteratively chosen to reduce model overfitting in highly sampled areas of the Estuary (Suisun Bay, Carquinez, etc.), while also providing acceptable spatial coverage in less intensively sampled areas.

The spatial mesh was geographically restricted to the wetted area of the Estuary by restricting spatial autocorrelation between geographically close, yet ecologically distinct, habitats (Figure 1). Shorelines were simplified using barrier polygons, and small channels were expanded to allow the mesh to fit an adequate number of knots for even representation of sparsely sampled areas. Likewise, only major islands separating heterogeneous habitats were included to increase the number of knots in the sinuous regions of the Delta and Suisun Marsh. Finally, the Montezuma Slough Salinity Control Gates and Delta Cross Channel Gates were treated as open to reflect the average condition during sampling periods (Figure 1). Simplification of the barrier polygon was an iterative process, as early barrier polygons with full shoreline and channel complexity had very few knots present in the Delta and Suisun Marsh regions.

The notational structure (Equation 1) of the binomial GLMMs is shown below. The number of sampling events that detected a given species at location *s* in year *t* and month *m* by sampling program *p, y*_*s,t,m,p*_ is modeled as a binomial random variable with expected probability of detection at a single sampling event equal to *μ*_*s,t,m,p*_ and *N*_*s,t,m,p*_ sampling events over the month. We used a logit link to model *μ*_*s,t,m,p*_ as a function of independent year effects (*α*_*t*_), survey effects (*β*_*p*_), and a cubic spline for month (*s(m)*). The variable *ω*_*s*_ represents spatial random effects, *ϵ*_*s,t*_ represents spatiotemporal random effects, and ζ_*t*_ represents the spatially varying coefficients through time *Y* (scaled year *t* centered around zero with a standard deviation equal to one). The random effects (*ω*_*s*_, *ϵ*_*s,t*_) and the spatially varying coefficient (*ζ*_*t*_) are drawn from Gaussian Markov random fields with Matérn covariance matrices Σ_*ω*_, Σ_*ϵ*_, and Σ_*ζ*_, respectively (Barnett et al. 2021). Spatial random effects were included to account for unmeasured variables that are approximately fixed through time (depth, distance upstream, substrate, etc.) whereas spatiotemporal random effects were included to account for unmeasured variables that are likely to change over both time and space (salinity, temperature, food availability, etc.; Anderson et al. 2022). Spatiotemporal random effects were treated as independent and identically distributed.

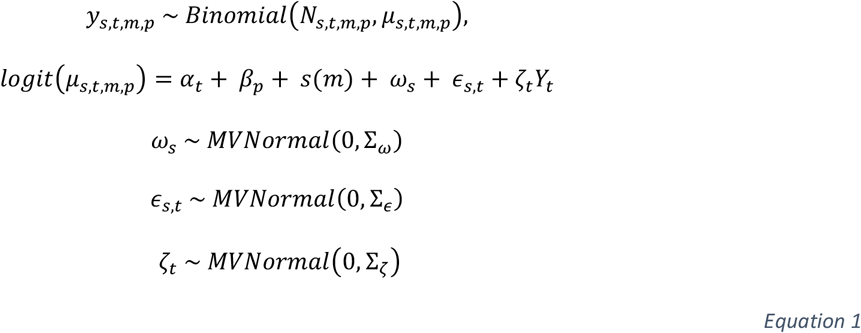

Residuals from the non-random effects were then simulated by drawing 500 samples from the fitted model and tested using the ‘DHARMa’ package in R (Hartig 2022). Residual uniformity was tested using the one-sample Kolmogorov-Smirnov test, residual dispersion was tested using the DHARMa nonparametric dispersion test via the standard deviation of residuals fitted versus simulated, and residual outliers were tested using the DHARMa outlier test based on the exact binomial test with approximate expectations (Hartig 2022). None of the models demonstrated unacceptable levels of residual uniformity, dispersion, or outliers, indicating good fit.

Previous publications have identified the potential pitfalls of generating models using disparate datasets in an integrated format (Walker et al. 2017; Moriarty et al. 2020; Huntsman et al. 2022). When unaccounted for, differences in survey effort, gear efficiency, and overall catchability can introduce significant biases in abundance and spatiotemporal density trends (Walker et al. 2017; Huntsman et al. 2022). However, our inclusion of survey as a fixed effect accounts for these biases, allowing separate intercepts to be fit for each of the eight surveys.

Once models were fit, predictions of the probability of detection and spatially explicit slopes of predictions over time were made for each of the five species across a 500m grid of the wetted area for the visualization of Estuary wide trends. Estimates of the spatial slope standard deviation as a metric of uncertainty were then generated by sampling (n=200) from the joint precision matrix.

Once prediction dataframes were made, the mean estimates of the probability of detection were calculated by decade for each spatial point and means were rasterized into smooth prediction planes using the R package ‘ggplot2’ (Wickham 2016). Rasterized spatial slopes, representing relative change through time, were plotted along with the standard deviation of the estimates of the spatial slopes. Mean annual estimates of the probability of detection at grid points for each of the five species were also plotted by applying a smoothing function (generalized additive model) by year.

To measure distributional sensitivity to changes in Delta outflow conditions (amount of water exiting the Delta after water exports) and overall shifts in population distributions, we calculated the annual predicted center of gravity (COG) and 95% confidence intervals along a longitudinal gradient for each of the modeled fish species. Delta outflow data was sourced from the California Department of Water Resources (CDWR 2022) as millions of acre-feet of water per calendar year. COG is a metric which represents the mean location of a population, weighted by density of observations, and although it is imperfect at describing local trends and/or detecting changes at distributional extremes (Barnett et al. 2021), it can be a useful metric for measuring population movement along a distributional gradient (Thorson et al. 2016). Due to computational limitations, COG estimates were generated using predictions at survey station points rather than at all points on the prediction grid. As a result, the COG of each species does not necessarily represent the true longitudinal center of each population, but changes in COG over time and relative differences between species are valid. COG was calculated longitudinally to best reflect the flow direction of the Estuary, which generally runs East-West from the Delta through the Central San Francisco Bay (Figure 1).

To test for the effects of Delta outflow on COG, temporal trends in COG, and differences in COG between species, we constructed a generalized additive model using the package “mgcv” (Pseudo-R Code, Equation 2; Wood 2011). The point estimates of COG for each species from the sdmTMB prediction outputs were included as the response variable, with yearly estimates weighted by their variance. Smooth functions with thin plate basis splines (bs=”tp”) were applied to “Year” by “Species” and “Delta Outflow” by “Species”, and “Species” was included as a linear fixed effect. Finally, generalized additive model results were printed in tabular format and plots were generated of yearly COG point estimates, 95% confidence intervals around the point estimates, the fit trendline and 95% confidence interval by species from the generalized additive model, and the spline effect of Delta outflow in million-acre feet on COG (Wickham 2016; Coretta 2022).

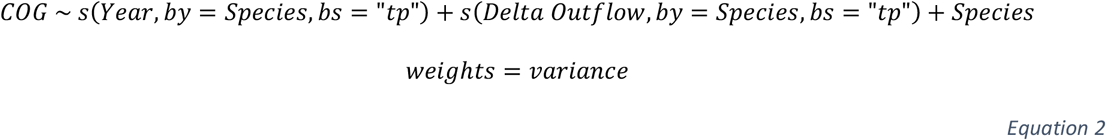

All analyses were conducted in R version 4.1.2 (R Core Team 2020). Code is available on github (https://github.com/dkstompe/SFE_Spatial_Fishes.git).

## Results

### Model Results

All models fully converged and fit the data acceptably well as determined through residual testing in the DHARMa package (Hartig 2022). The results of residual dispersion tests and residual outlier tests were significant for some models (Appendix Table 1); however, this was driven by the exceptionally large number of data points included in the model rather than poor model fit (Hartig 2022). For example, the dispersion value of 0.992 for the American shad model was significant (p=0.012) as was the outlier test (p=0.0083) with just 467 outliers out of 103,341 observations. The results of model diagnostics are included in the supplementary material (Appendix Figure 2, Appendix Table 1).

**Figure 2.**
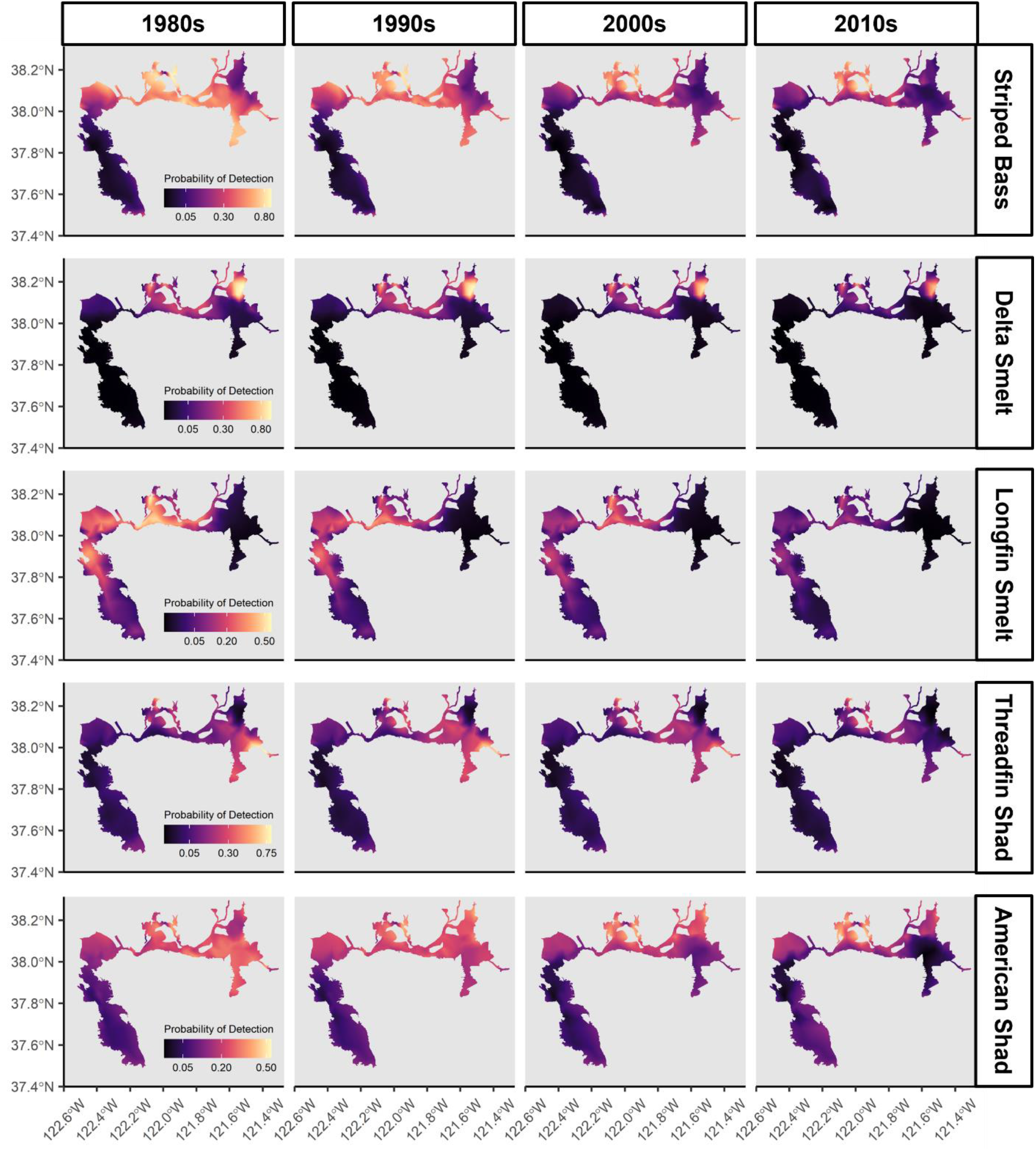
Mean probability of distribution of striped bass, Delta smelt, longfin smelt, threadfin shad, and American shad by decade, as predicted by GLMMs. Hotter colors denote higher probability of detection, cooler colors lower probability of detection. Note: color scales are on a square root scale.

The output of the GLMMs showed differing coefficients by ‘Survey’ for all species as well as negative coefficients for the linear component of the smooth effect of ‘Month’ sampled for threadfin shad and American shad (Table 2). The differing coefficients by ‘Survey’ indicate differences in the catchability of species by survey methodology and gear type, while the negative coefficient of the linear component of ‘Month’ is likely due to seasonal differences in either local abundance, absolute abundance, or vulnerability of capture across life stages for the two shad species.

**Table 2.** Model results from binomial generalized linear mixed models of probability of detection for striped bass, Delta smelt, longfin smelt, threadfin shad, and American shad. Table contains factor (Survey) and smooth (Month) model coefficients and standard errors. The Intercept term is assigned to the CDFW Bay Study Midwater Trawl. Matérn range is the distance at which spatial correlation degrades to ∼0.13. Survey abbreviations are as follows: BOT = CDFW Bay Study Otter Trawl, BSS = USFWS Beach Seine Survey, CIT = USFWS Chipps Island Trawl, FMWT = CDFW Fall Midwater Trawl, SMBS = UC Davis Suisun Marsh Beach Seine, SMOT = UC Davis Suisun Marsh Otter Trawl, STN = CDFW Summer Townet Survey.

|  | Striped Bass |  | Delta Smelt |  | Longfin Smelt |  | Threadfin Shad |  | American Shad |  |
| --- | --- | --- | --- | --- | --- | --- | --- | --- | --- | --- |
|  | coef.est | coef.se | coef.est | coef.se | coef.est | coef.se | coef.est | coef.se | coef.est | coef.se |
| **(Intercept)** | 0.79 | 0.41 | -4.89 | 0.92 | -4.16 | 0.51 | -1.72 | 0.82 | -1.68 | 0.29 |
| **Survey:BOT** | -0.09 | 0.04 | -1.41 | 0.08 | 0.29 | 0.03 | -1.77 | 0.1 | -2.56 | 0.07 |
| **Survey:BSS** | -2.70 | 0.09 | -2.50 | 0.15 | -3.8 | 0.29 | -0.27 | 0.1 | -3.32 | 0.12 |
| **Survey:CIT** | -0.40 | 0.07 | 0.96 | 0.10 | -0.84 | 0.08 | 0.63 | 0.13 | 2.51 | 0.09 |
| **Survey:FMWT** | -0.29 | 0.04 | 0.14 | 0.07 | 0.16 | 0.05 | 0.27 | 0.07 | 0.14 | 0.05 |
| **Survey:SMBS** | -1.20 | 0.17 | -1.76 | 0.44 | -1.95 | 0.32 | -0.52 | 0.25 | -3.01 | 0.27 |
| **Survey:SMOT** | -0.08 | 0.12 | -2.31 | 0.26 | -0.34 | 0.16 | -1.86 | 0.19 | -2.95 | 0.17 |
| **Survey:STN** | -0.61 | 0.05 | 0.87 | 0.08 | -0.54 | 0.06 | -0.69 | 0.1 | -1.67 | 0.07 |
| **s(Month)** | -0.04 | 0.00 | 0.00 | 0.01 | -0.01 | 0 | -0.19 | 0.01 | -0.26 | 0.01 |
| **Matern Range** | 29.69 |  | 29.20 |  | 27.30 |  | 36.52 |  | 24.47 |  |

As expected, the pelagic gear types used by surveys such as the Bay Study Midwater Trawl, the Fall Midwater Trawl, and the Chipps Island Trawl were generally most effective at catching the pelagic species included in our analysis as indicated by model coefficients, with some notable exceptions (Table 2). For example, the Fall Midwater Trawl, a survey specifically designed to capture young of year striped bass, had a lower model coefficient than the reference survey (Bay Study Midwater Trawl). Beach seines and benthic trawls typically had negative coefficients amongst the modeled fish species, indicating lower capture efficiencies than the reference survey (Bay Study Midwater Trawl).

### Spatial Trends

In general, species show an overall reduction in spatial probabilities of detection over the modeled time period (Figure 2). Spatial slopes are negative in most regions for most species and are exclusively negative or near zero for Delta smelt and longfin smelt (Figure 3). Positive slope values are present in limited regions for striped bass (Suisun Marsh, North Delta, South San Francisco Bay), threadfin shad (Confluence, Suisun Bay, Suisun Marsh), and American shad (North Delta, Suisun Bay, Suisun Marsh, South San Francisco Bay).

**Figure 3.**
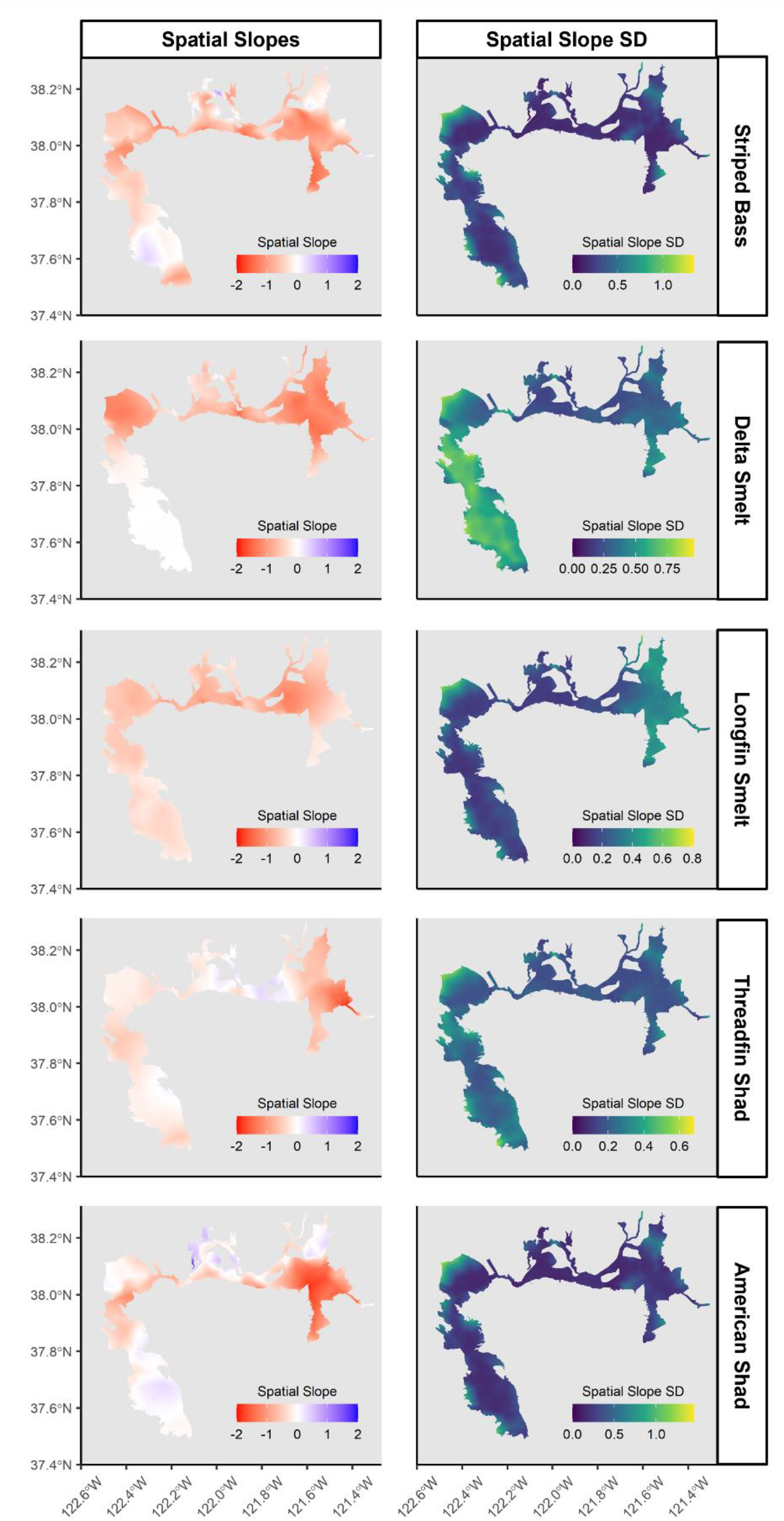
Spatial slopes and standard deviations (SD) of spatial slopes for the five modeled fish species. Red slope shading indicates a decrease in the probability of detection between 1980 and 2017, white is no change, and blue indicates increased probability of detection. Hotter colors indicate higher SD and thus increased uncertainty in model predictions of spatial slope.

Species show some shared patterns in distributional changes in the probability of detection over time; most notable of which is the reduction in estimates in the Central and South Delta (Figure 1 – regions 8-9; Figure 2). This trend exists for striped bass, threadfin shad, and American shad which historically had relatively high probabilities of detection in these regions, but which had very low detection probabilities by the 2010s. Further supporting this are the spatial slopes which are strongly negative in the Central and South Delta for these species (Figure 3). The reduction in detection probability in these regions drives a constriction in distribution away from large parts of the Delta and towards the Confluence and Suisun regions (Figure 1 – region 6 and 5).

Unlike the other three species, the two smelt species were rarely detected in the Central and South Delta regions at any point during the modeled time period. Longfin smelt were mostly distributed downstream, with historically high probabilities of detection in the Central San Francisco Bay through the Confluence region (Figure 1 -regions 2-5). Conversely, Delta smelt were not found in the lower portions of the Estuary and were instead most likely to be detected in the North Delta, Confluence, and Suisun Regions (Figure 1 – regions 5-7). Both species seem to exhibit a reduction in overall detection probabilities rather than a constriction of distribution.

Another notable trend is the persistently higher probability of detection in the Suisun Marsh and Suisun Bay regions (Figure 1 – region 5 top and bottom, respectively) relative to other parts of the Estuary for all species. For most species it appears that the probability of detection does not increase in Suisun over the decades, but rather it decreases less. This is supported by the spatial slope plots which show relatively less negative slope in the Suisun Bay and especially Suisun Marsh regions (Figure 3). American shad was the one species which did have positive slopes throughout much of the Suisun Marsh region, indicating an increased detection probability between 1980 and 2017.

In general, there was little uncertainty in the spatial slopes in highly sampled regions where species were detected by the eight surveys over the modeled time period. Uncertainty was high at the edge of the spatial mesh, such as the north end of San Pablo Bay (Figure 1 – region 3), or where species were never or rarely detected (Delta Smelt, Central and South San Francisco Bays, Figure 2 & 3). Spatial slopes should be interpreted cautiously in these specific areas given the relatively high level of uncertainty.

### Detection Trends

The overall probability of detection for the included species generally declined between 1980 and 2017 (Figure 4), evidence that abundance may be declining. Declines are most evident for striped bass and the two smelt species, the latter of which are now rarely detected in the eight surveys. Conversely, the detection probabilities for striped bass, threadfin shad, and American shad appear to somewhat rebound near 2017 after lows in 2010 through 2012.

**Figure 4.**
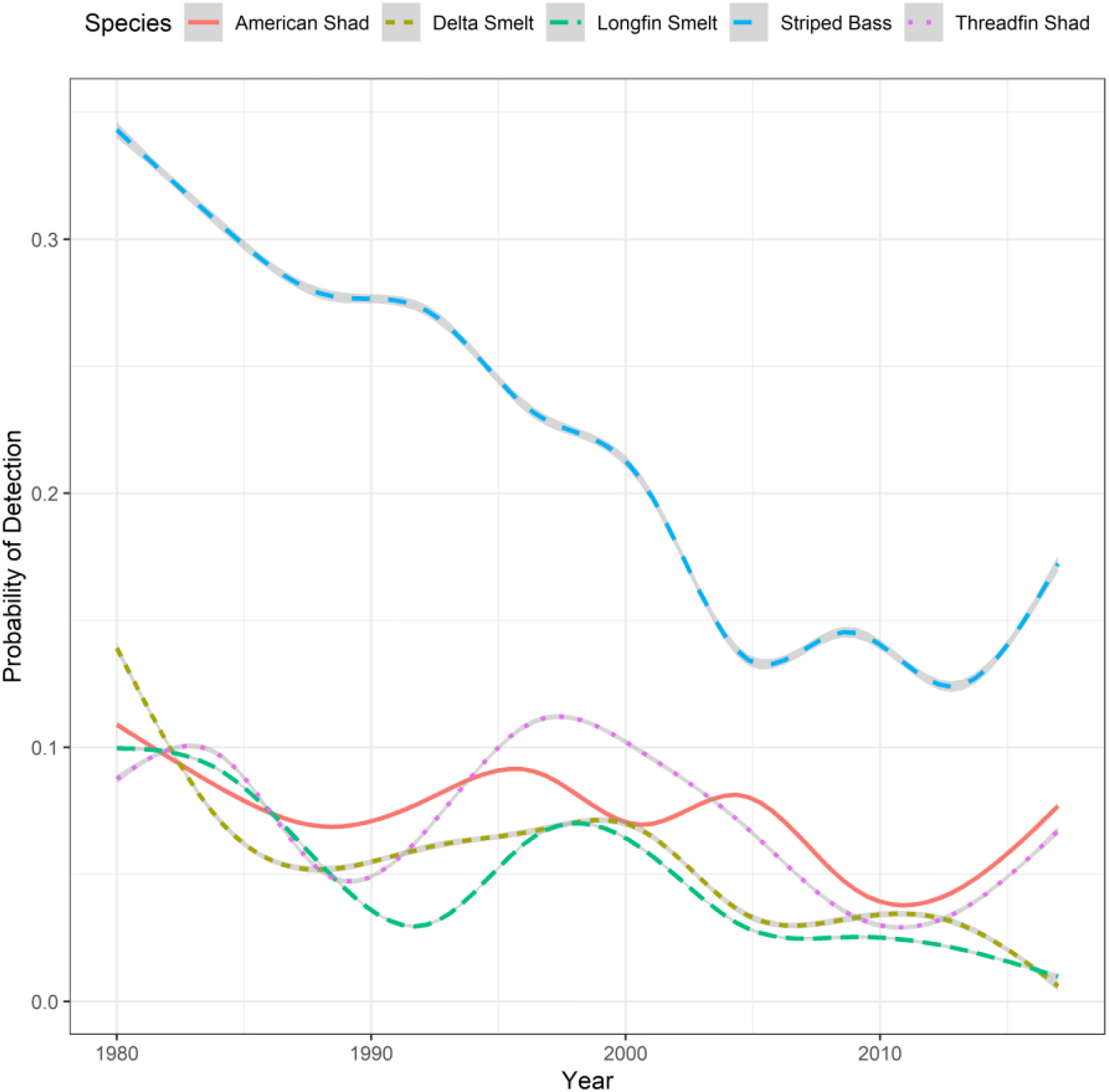
Overall trends in the predicted probability of detection by the eight-survey aggregate dataset as calculated by generalized additive model smoother of estimates at 500m grid points.

The trends in detection probability are primarily nonlinear, with intermittent periods of increase or stabilization. Striped bass have the largest overall decline, from a detection probability of approximately 0.35 in the early 1980s to less than 0.15 by 2012 (Figure 4). Delta smelt have an initial steep decline in the early 1980s, followed by relatively stable to increasing detection probability until another period of steep decline in the early 2000s. After this decline Delta smelt again stabilize until the mid-2010s, at which point they decline to near zero. Longfin smelt trends are similar to Delta smelt, but with a less dramatic initial decline followed by a low in the early 1990s and a rebound in the late 1990s before ultimately declining to near zero as well. Threadfin shad trends are somewhat similar to longfin smelt, but as stated earlier, they have partially recovered in the years since 2010. Finally, American shad are unique in their trends, with a somewhat stable detection probability until a decline in the late 2000s, followed by a recovery after 2010.

### Center of Gravity

The generalized additive model (Equation 2) of the smooth effects of year and Delta outflow by species on COG, and of the linear effects of species on COG, fit with an adjusted R squared of 0.969 and explained 97.7% of the deviance in the data.

The center of gravity (COG) of each fish species partitioned by easting, with threadfin shad distributed the furthest upstream (Table 3, Figure 5). This is followed by Delta smelt, American shad, and striped bass clustered within approximately 5km of one another, and longfin smelt the furthest downstream (Table 3, Figure 5).

**Table 3.**
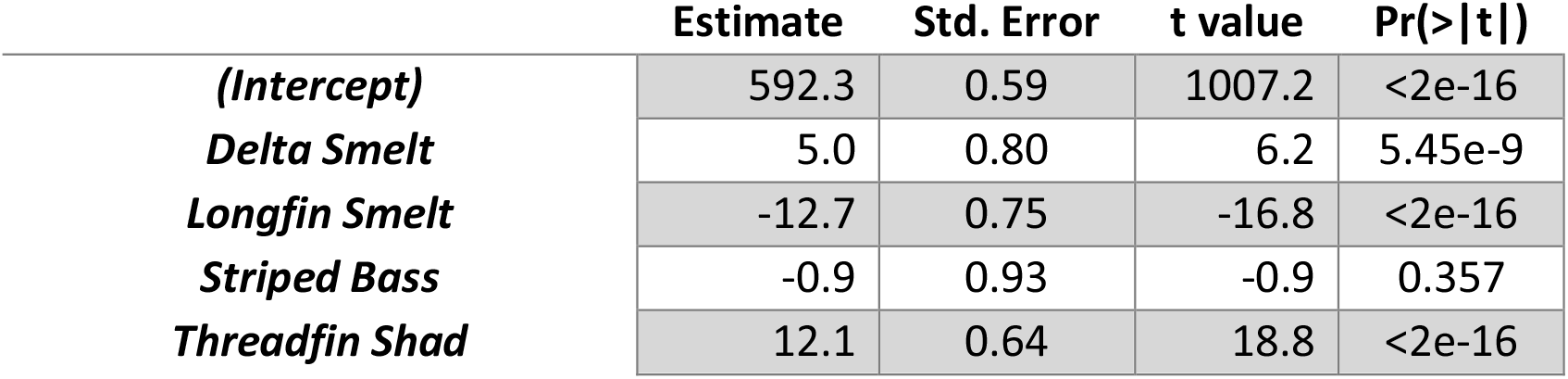
Model results for the parametric linear terms of COG generalized additive model. Intercept represented by American shad.

**Figure 5.**
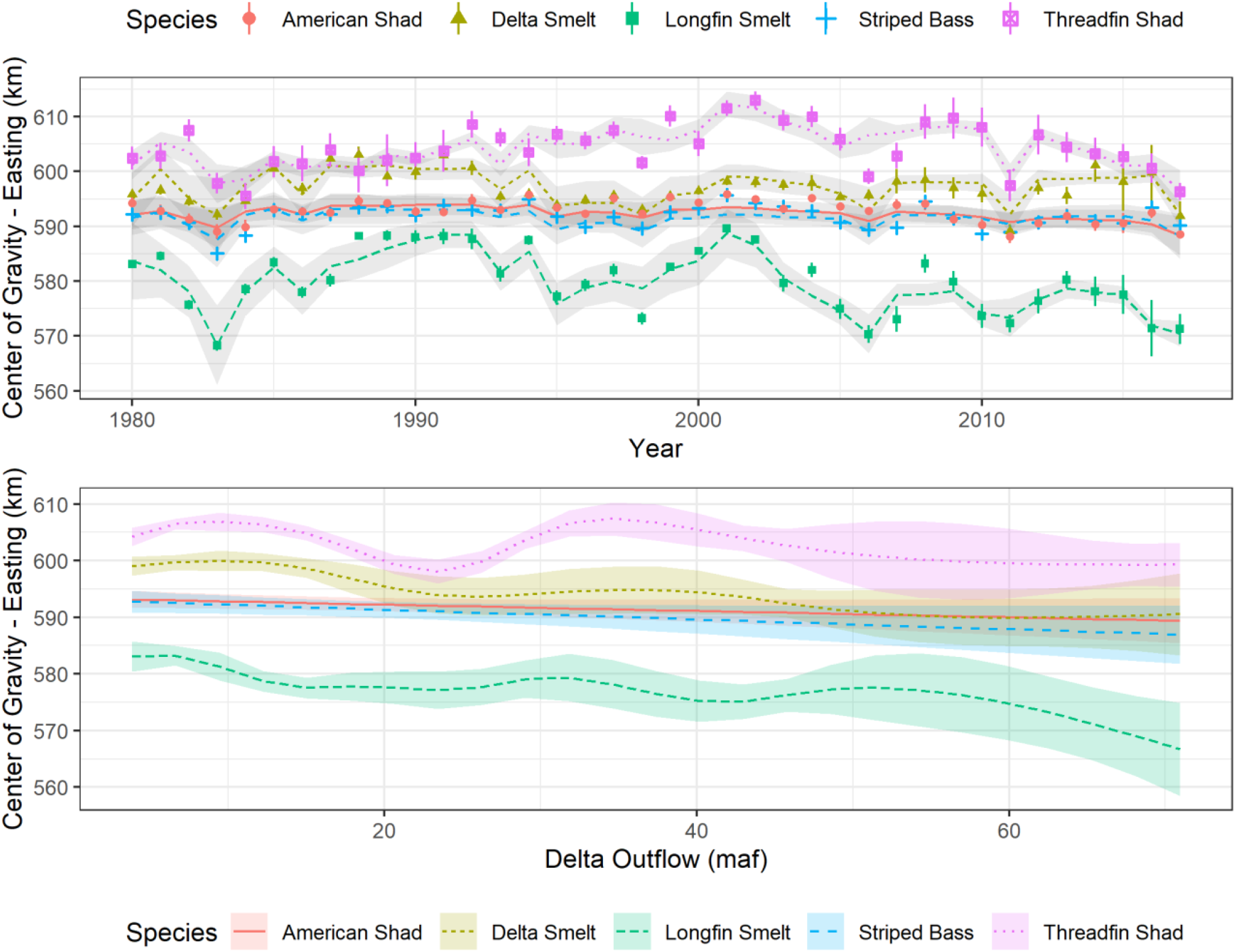
Top panel is the center of gravity (COG) of the five modeled fish species from 1980-2017, shown as yearly point estimates with 95% confidence intervals as well as via generalized additive model fit (Equation 2). Bottom panel are the estimated smooths of the center of gravity for each species across values of Delta outflow, measured in million-acre feet (maf).

The smooth term for Delta outflow indicated effects of Delta outflow on COG for longfin smelt (p=1.63e-6), Delta smelt (p=6.92e-7), and threadfin shad (p<2e-16), while little to no effect was seen for striped bass (p=0.069) or American shad (p=0.123; Table 4). While effects clearly existed for Delta smelt, longfin smelt, and threadfin shad, the magnitude and direction of shifts in COG at different Delta outflow values were not uniform amongst the species (Figure 5). Longfin smelt were the most affected by changes in outflow conditions, with a somewhat linear relationship and an estimated center of gravity more than 15km downstream at high Delta outflow than at the lowest outflow values (Figure 5). Delta smelt COG also declined somewhat linearly as Delta outflow increased, although at a lower magnitude than for longfin smelt (Figure 5). Changes in threadfin shad COG shifted considerably under different outflow conditions; however, this relationship was non-linear with minima at both 23 and 60+ maf (Figure 5).

**Table 4.** Model results for the smooth interaction terms from the COG generalized additive model.

|  | edf | Ref.df | F | p-value |
| --- | --- | --- | --- | --- |
| ***s*(Year):American Shad** | 1.78 | 2.21 | 1.99 | 0.132 |
| ***s*(Year):Delta Smelt** | 2.87 | 3.49 | 2.40 | 0.071 |
| ***s*(Year):Longfin Smelt** | 8.50 | 8.90 | 13.59 | <2e-16 |
| ***s*(Year):Striped Bass** | 1.00 | 1.00 | 0.51 | 0.477 |
| ***s*(Year):Threadfin Shad** | 8.60 | 8.94 | 13.14 | <2e-16 |
| ***s*(Delta Outflow):American Shad** | 1.00 | 1.00 | 2.41 | 0.123 |
| ***s*(Delta Outflow):Delta Smelt** | 5.09 | 5.98 | 7.73 | 6.92e-7 |
| ***s*(Delta Outflow):Longfin Smelt** | 6.88 | 7.83 | 6.01 | 1.63e-6 |
| ***s*(Delta Outflow):Striped Bass** | 1.00 | 1.00 | 3.37 | 0.069 |
| ***s*(Delta Outflow):Threadfin Shad** | 7.67 | 8.48 | 9.26 | <2e-16 |

The point estimates and modeled fit of COG by species also resulted in several distinct patterns over the modeled time period. The smooth term for year indicated effects on COG (Table 4) for longfin smelt (p<2e-16) and threadfin shad (p<2e-16), little effect for Delta smelt (p<0.071), and no effect for American shad (p=0.132) or striped bass (p=0.477). Longfin smelt had the most dramatic temporal trends in COG, spanning approximately 20km over the modeled time period (Figure 5). In addition, longfin smelt COGs remained further downstream during the period after 2002, despite some periods of extreme drought (CDEC 2022). Threadfin shad also showed strong temporal trends in COG, but with a unique pattern where the population generally shifted upstream during the period from approximately 1990 through 2010 (Figure 5). Finally, Delta smelt remained relatively centered within a 5km band, with little interannual variability (Figure 5).

## Discussion

Using new developments in spatiotemporal modeling, we leveraged the rich but fragmented monitoring data in the Estuary to demonstrate changes in the spatial and temporal probability of detecting key pelagic fish species during a period of considerable abiotic and biotic change. Large swaths of the Estuary that historically supported high detection probabilities of striped bass, threadfin shad, and American shad, including the South and Central Delta, are now relatively devoid of these species, driving population constrictions to the Suisun and Confluence regions. Over the same time period, Delta smelt and longfin smelt experienced relatively even declines throughout the Estuary. The detection probability of all species declined to some extent from their levels in 1980; however, the trends and future outlooks differ by species.

It is important to note that differences in life history and age-structured distribution of the species can color interpretation. Striped bass is a long-lived semi-anadromous species most likely to be caught by survey gear during their first year of life; results reflect juvenile distribution and are not directly indicative of adult behavior or abundance. Likewise, American shad typically migrate out of the Estuary (and into the Pacific Ocean) after their first year (Carothers et al. 2021), so juveniles are best represented in our results. Longfin smelt also sometimes leave the Estuary; however, they are susceptible to survey gear throughout their lives so results may be interpreted as representing the total local population. Finally, Delta smelt and threadfin shad primarily remain within the Estuary and are vulnerable to survey gear throughout their short (1-3) year lifespans. Specifically, Delta smelt is an annual species (Moyle 2002) fully restricted to the sampled estuarine areas so our results may be interpreted as representing the spatiotemporal trends of the species as a whole.

Given these species-specific differences in model interpretation, it is clear that Delta smelt and longfin smelt have declined precipitously since the 1980s (Figure 4). These species are now rarely detected by Estuary surveys. The threadfin shad population has also experienced declines in overall abundance; however, it appears to be more robust in its ability to shift to different regions as evidenced by positive spatial slopes in the Confluence and Suisun regions (Figure 3). As a result, threadfin shad overall probability of detection has somewhat recovered since 2010 (Figure 4). The most dramatic decline in detection probability is seen for striped bass (Figure 4), indicating a reduction in either spawning success or juvenile survival over the modeled time period. American shad spawning success or juvenile survival has also somewhat declined between 1980 and 2017, although their probability of detection is now only slightly below historic highs after a recovery since 2010 (Figure 4). The lower level of overall decline in juvenile American shad versus juvenile striped bass, despite similar distributional patterns, may be somewhat driven by increased utilization of Suisun Marsh by American shad as the Central and South Delta became inhospitable (Figure 3).

Sensitivity of annual COGs to outflow conditions indicate different relative effects of climate and water management for each of the species. Delta outflow has major effects on the location of the salinity gradient, and thus the highly productive low-salinity zone (MacWilliams et al. 2015). Given that longfin smelt COGs are associated with Delta Outflow (Table 4, Figure 5), this indicates that the distribution of this species may annually shift to track areas of favorable salinity and/or productivity. Conversely, species such as American shad whose COGs are insensitive to different outflow conditions (Table 4, Figure 5) may not be as plastic in their annual distribution, potentially due to reliance on habitat structure rather than water conditions. It is difficult to identify whether longitudinal plasticity in response to changes in Delta outflow is advantageous, as both species with highly variable COGs, such as longfin smelt, and species with relatively stable COGs, such as striped bass, have both experienced dramatic declines in their probability of detection.

Trends in annual COG over time suggest differential responses to changing environmental conditions and highlight life history differences among the species (Table 3, Figure 5). The highly variable annual COGs for longfin smelt and threadfin shad support a plastic response in distribution, or potentially differences in success between multiple subpopulations within the Estuary. For example, multiple spawning populations of longfin smelt have been identified within the Estuary (Lewis et al. 2019), so regional differences in spawning success or survival could shift annual COGs. Conversely, the relatively fixed annual COGs for Delta smelt, striped bass, and American shad indicate a general reliance on fixed habitat features and either no subpopulation structure or subpopulations with similar interannual spawning success or survival.

The abiotic and biotic drivers of the described changes in abundance and distribution are likely complex and interacting. For example, over the modeled time period, the Estuary saw changes in water export regimes (Gartrell et al. 2017), the introduction of several highly invasive plant and invertebrate species (Cohen and Carlton 1998), and both record-setting droughts (Durand et al. 2020) and extremely wet years (CDEC 2021). These factors interact, changing the amount and quality of habitat for native and introduced pelagic fishes.

For example, invasive plants such as Brazilian waterweed (*Egeria densa*) have benefitted from reduced turbidity due to upstream impoundments and the constant freshwater condition maintained by water export operations (Durand et al. 2016). Dense stands of Brazilian waterweed have reduced water velocity in some areas, dropping out additional suspended particulate matter and capturing nutrients from upstream sources (Yarrow et al. 2009; Durand et al. 2016). This has resulted in potentially reduced pelagic productivity (Vanderstukken et al. 2011; Durand et al. 2016), a shift in zooplankton communities important for small and larval fish diets (Espinosa-Rodríguez et al. 2021), and reduced turbidity, which makes small fishes more susceptible to predation (Ferrari et al. 2014). There are myriad examples of such interacting and cascading effects that have reduced the suitability of the pelagic habitat within the Estuary (Brown and Moyle 2005; Sommer et al. 2007; Brooks et al. 2012; Cloern and Jassby 2012; Sabal et al. 2016).

An overarching trend in the distribution and regional abundance of these species is the relative insulation of the Suisun Region, and to a lesser degree the North Delta, from overall declines in detection probability (Figure 2, 3). There are many potential drivers of this, one of which is likely the historically lower levels of colonization of submersed aquatic vegetation, such as Brazilian Waterweed, in these regions. Brazilian waterweed is largely limited by salinity in Suisun Marsh and Bay (Borgnis and Boyer 2016) and was previously limited by turbidity in the North Delta (Durand et al. 2016). Given their relatively lower levels of decline in detection probability, these regions could prove important for maintaining viable populations of pelagic fishes in the future.

Despite lower levels of decline or even increased detection probability in the Suisun region and the North Delta, it should be noted that these regions may not necessarily represent *ideal* habitat in the face of system wide degradation. These regions may simply be *better* than those regions which have experienced substantial declines in detection probability (Central Delta, South Delta). Under this scenario, fish may be shunted away from previously productive habitats into regions which have experienced relatively less change. This likely partially explains the increased detection probability of some species in Suisun Bay and Marsh and the North Delta over the modeled time period.

It should be noted that the station density is somewhat sparse in the North and South Delta, with one or few surveys representing catch in these regions. Specifically, trends in the North Delta are most influenced by catch from the USFWS Beach Seine Survey, while trends in the south Delta are most influenced by catch from the Summer Townet Survey (Appendix Figure 1). These regional differences in station density and representation should be considered when interpretating the absolute detection probability of species such as Delta smelt in the North Delta.

## Conclusions

By leveraging existing long-term survey data in an integrated modeling format, we have described trends in the distribution and abundance of five key Estuary fish species. The modeling techniques we employ have most commonly been used to describe trends in large adult marine fishes, but we demonstrate their ability to model trends in juvenile or small estuarine fishes as well. Our approach also demonstrates the value of using an integrated data set due to the greatly increased spatial density and coverage. These data can detect distributional trends that would otherwise not be covered by a single survey. We are aware that measures must be taken to ensure that modeling with disparate data does not impart unacceptable biases, so we included ‘survey’ as a fixed effect and only used consistently surveyed stations. These should be sufficient to control for the methodological disparities. Our modeling and data integration methods should not only prove useful for management of Estuary fishes, but also for describing trends in distribution and abundance of fishes in other inland, estuarine, and marine systems with multiple independent surveys.

The individual long-term fish surveys of the Estuary have collected valuable data for tracking trends in the distribution and abundance of the species we considered here, and indeed for most fish species found within the Estuary. While any one survey can describe part of a species’ story, it is only when surveys are analyzed in concert that we can see the true extent of change. The increased spatial breadth and detail of an integrated analysis allows us to see much more granular and localized changes in distribution.

The Estuary has experienced dramatic changes to its hydrology, biotic communities, and physical structure, which in turn has reduced the detection probability and distributional breadth of both native and naturalized pelagic fish species. The species analyzed here include fish that hold considerable ecological, recreational, and cultural value amongst California stakeholders. We show, through distributional shifts and spatial slopes, that a major driver in the reduced detection probability of pelagic fishes is the declines in their populations in large portions of the Delta. Conversely, Suisun Marsh and the North Delta appear to function as refuge habitats for at least a few of the species, reinforcing that they should be managed as high priority refuges for conservation.

While our analyses identify regions and time periods of change for these fish species, they do not specifically identify biological or abiotic drivers of these changes. In future efforts, our models may be expanded through the inclusion of spatially explicit data for both biotic and abiotic predictors, including bathymetry, temperature, and LIDAR imagery of submersed aquatic vegetation. Expansion of our models in this way would further refine our understanding of important habitat criteria for pelagic fishes in the Estuary, which should result in more effective habitat restoration and conservation.

## Acknowledgements

Caroline Newell and Dr. Eric Ward provided helpful reviews of an early version of this manuscript. This study would not have been possible without the numerous staff and managers of Estuary fish surveys who spent considerable time and effort collecting the data that we used. This project was funded in part by the California Department of Fish and Wildlife and the California Department of Water Resources.

## Notes

https://github.com/dkstompe/SFE_Spatial_Fishes.git

### Competing Interest Statement

The authors have declared no competing interest.

## References

Anderson, M.G. 2005. Habitat restoration in the Columbia River Estuary: A strategy for implementing standard monitoring protocols. Thesis. Oregon State University, Corvallis, Oregon, USA.

Anderson, S.C., E.A. Keppel, and A.M. Edwards. 2019. A reproducible data synopsis for over 100 species of British Columbia groundfish. Report 2019/041. Canadian Science Advisory Secretariat, Fisheries and Oceans Canada, Ottawa, ON.

Anderson, S.C., E.J. Ward, P.A. English, and L.A.K. Barnett. 2022. sdmTMB: an R package for fast, flexible, and user-friendly generalized linear mixed effects models with spatial and spatiotemporal random fields. Preprint. bioRxiv. https://doi.org/10.1101/2022.03.24.485545.

Barnett, L.A.K., E.J. Ward, and S.C. Anderson. 2021. Improving estimates of species distribution change by incorporating local trends. Ecography 44(3): 427–439.

Baxter, R., K. Hieb, S. DeLeón, K. Fleming, and J. Orsi. 1999. Report on the 1980-1995 Fish, Shrimp, and Crab Sampling in the San Francisco Estuary. Technical Report 53. Sacramento, CA: California Department of Fish and Game.

Blaber, S.J.M., K.W. Able, and P.D. Cowley. 2022. Estuarine fisheries. In Fish and fishes in estuaries: A global perspective, ed A.K. Whitfield, K.W. Able, S.J.M. Blaber, and M. Elliott, 553–616. Hoboken: Wiley.

Brooks, M.L., E. Fleishman, L.R. Brown, P.W. Lehman, I. Werner, N. Scholz, C. Mitchelmore, J.R. Lovvorn, M.L. Johnson, D. Schlenk, S. van Drunick, J.I. Drever, D.M. Stoms, A.E. Parker, and R. Dugdale. 2012. Life histories, salinity zones, and sublethal contributions of contaminants to pelagic fish declines illustrated with a case study of San Francisco Estuary, California, USA. Estuaries and Coasts 35(2): 603–621.

Brown, L.R., and P.B. Moyle. 2005. Native fishes of the Sacramento–San Joaquin Drainage, California: A history of decline. American Fisheries Society Symposium 45: 75–98.

Borgnis, E., and K.E. Boyer. 2016. Salinity tolerance and competition drive distributions of native and invasive submerged aquatic vegetation in the Upper San Francisco Estuary. Estuaries and Coasts 39(3): 707–717.

Cabral H.N., A. Borja, V.F. Fonseca, T.D. Harrison, N. Teichert, M. Lepage, and M.C. Leal. 2022. Fishes and estuarine environmental health. In Fish and fishes in estuaries: A global perspective, ed A.K. Whitfield, K.W. Able, S.J.M. Blaber, and M. Elliott, 332–379. Hoboken: Wiley.

Carothers, C., J. Epifanio, S. Gregory, D. Infante, W. Jaeger, C. Jones, P.B. Moyle, T.P. Quinn, K. Rose, T. Turner, T. Wainwright. 2021. American Shad in the Columbia River: Past, present, future. Report 2021-4. Independent Scientific Advisory Board, Northwest Power and Conservation Council, Portland, OR.

CDEC. 2021. Chronological reconstructed Sacramento and San Joaquin Valley water year hydrologic classification indices. California Department of Water Resources. Available from: https://cdec.water.ca.gov/reportapp/javareports?name=WSIHIST

CDWR. 2022. Dayflow. Suisun Marsh Branch, California Department of Water Resources. Available from: data.cnra.ca.gov/dataset/dayflow

Chadwick, H.K., D.E. Stevens, and L.W. Miller. 1977. Some factors regulating the striped bass population in the Sacramento–San Joaquin Estuary, California. Proceedings of the Conference on assessing the effects of power-plant-induced mortality on fish populations, ed W.V. Winkle, 18–35. Oxford: Pergamon Press Inc.

Cloern, J.E. and A.D. Jassby. 2012. Drivers of change in estuarine-coastal ecosystems: Discoveries from four decades of study in San Francisco Bay. Reviews of Geophysics 50(4).

Cohen, A.N., and J.T. Carlton. 1998. Accelerating invasion rate in a highly invaded estuary. Science 279(5350): 555–558.

Coretta, S. 2022. tidymv: tidy model visualisation for generalised additive models. R package version 3.3.0. https://CRAN.R-project.org/package=tidymv

Cowley, P.D., J.R. Tweedley, and A.K. Whitfield. 2022. Conservation of estuarine fishes. In Fish and fishes in estuaries: A global perspective, ed A.K. Whitfield, K.W. Able, S.J.M. Blaber, and M. Elliott, 617-683. Hoboken: Wiley.

Dekar M.P., P.L. Brandes, J. Kirsch, L. Smith, J. Speegle, P. Cadrett, and M. Marshall. 2013. Background report prepared for review by the IEP Science Advisory Group, June 2013. Lodi, CA: U.S. Fish and Wildlife Service.

Dill, W.A. and A.J. Cardone. 1997. History and status of introduced fishes in California, 1871 – 1996. California Department of Fish and Game. Fish Bulletin 178.

Durand, J., F. Bombardelli, W. Fleenor, Y. Henneberry, J. Herman, C. Jeffres, M. Leinfelder-Miles, R. Lusardi, A. Manfree, J. Medellín-Azura, B. Milligan, P. Moyle, and J. Lund. 2020. Drought and the Sacramento–San Joaquin Delta, 2012–2016: Environmental Review and Lessons. San Francisco Estuary and Watershed Science 18(2).

Durand, J., W. Fleenor, R. McElreath, M.J. Santos, and P. Moyle. 2016. Physical controls on the distribution of the submersed aquatic weed Egeria densa in the Sacramento–San Joaquin Delta and implications for habitat restoration. San Francisco Estuary and Watershed Science 14(4).

Espinosa-Rodríguez, C.A., S.S.S. Sarma, and S. Nandini. 2021. Zooplankton community changes in relation to different macrophyte species: Effects of Egeria densa removal. Ecohydrology & Hydrobiology 21(1): 153–163.

Ferrari, M.C.O., L. Ranåker, K.L. Weinersmith, M.J. Young, A. Sih, and J.L. Conrad. 2014. Effects of turbidity and an invasive waterweed on predation by introduced largemouth bass. Environmental Biology of Fishes 97(1): 79–90.

Ferguson, E. 2016. Trends in angling effort, catch, and harvest of American Shad, and implications for regulations in the Sacramento Basin sport fishery. Memorandum. West Sacramento: California Department of Fish and Wildlife.

Feyrer, F., T. Sommer, and S.B. Slater. 2009. Old school vs. new school: Status of Threadfin Shad (Dorosoma petenense) five decades after its introduction to the Sacramento–San Joaquin Delta. San Francisco Estuary and Watershed Science 7(1).

Gartrell, G., J. Mount, E. Hanak, and B. Gray. 2017. A new approach to accounting for environmental water: insights from the Sacramento–San Joaquin Delta. Report. Sacramento, CA: Public Policy Institute of California.

Godø, O.R., S.J. Walsh, and A. Engås. 1999. Investigating density-dependent catchability in bottom-trawl surveys. ICES Journal of Marine Science 56(3): 292–298.

Hartig, F. 2022. DHARMa: Residual diagnostics for hierarchical (multi-Level / mixed) regression models. R package version 0.4.5. https://CRAN.R-project.org/package=DHARMa

Herbold, B., D.M. Baltz, L. Brown, R. Grossinger, W. Kimmerer, P. Lehman, P.B. Moyle, M. Nobriga, and C.A. Simenstad. 2014. The role of tidal marsh restoration in fish management in the San Francisco Estuary. San Francisco Estuary and Watershed Science 12(1).

Herrgesell, P.L. 2012. A historical perspective of the Interagency Ecological Program: Bridging multi-agency studies into ecological understanding of the Sacramento-San Joaquin Delta and Estuary for 40 Years. Report. Sacramento, CA: California Department of Fish and Game.

Honey, K., R. Baxter, Z. Hymanson, T. Sommer, M. Gingras, and P. Cadrett. 2004. IEP long-term fish monitoring program element review. Interagency Ecological Program for the San Francisco Bay/Delta Estuary. Available from: https://www.academia.edu/507408/IEP_long_term_fish_monitoring_program_element_re-view

Huntsman, B., B. Majardja, and S. Bashevkin. 2022. Relative bias in catch among long-term fish monitoring surveys within the San Francisco Estuary. San Francisco Estuary and Watershed Science 20(1).

Malinich, T.D. 2020. Summer townet survey. 2020 Factsheet. Sacramento, CA: Interagency Ecological Program. Available from: https://nrm.dfg.ca.gov/FileHandler.ashx?DocumentID=185025

Kohlhorst, D.W. 1999. Status of striped bass in the Sacramento-San Joaquin estuary. California Fish and Game 85(1): 31–36.

Lewis, L.S., M. Willmes, A. Barros, P.K. Crain, and J.A. Hobbs. 2020. Newly discovered spawning and recruitment of threatened longfin smelt in restored and underexplored tidal wetlands. Ecology 101(1).

Lotze, H.K., H.S. Lenihan, B.J. Bourque, R.H. Bradbury, R.G. Cooke, M.C. Kay, S.M. Kidwell, M.X. Kirby, C.H. Peterson, and J.B. Jackson. 2006. Depletion, degradation, and recovery potential of estuaries and coastal seas. Science 312(5781):1806–1809.

MacWilliams, M.L., A.J. Bever, E.S. Gross, G.S. Ketefian, and W.J. Kimmerer. 2015. Three-dimensional modeling of hydrodynamics and salinity in the San Francisco Estuary: An evaluation of model accuracy, X2, and the low-salinity zone. San Francisco Estuary and Watershed Science 13(1).

McKenzie, R. 2021a. 2019 Delta juvenile fish monitoring program. Salmonid annual report. Lodi, CA: US Fish and Wildlife Service.

McKenzie, R. 2021b. 2019–2020 Delta juvenile fish monitoring program. Nearshore fishes annual report. Lodi, CA: US Fish and Wildlife Service.

Moriarty, M., D. Pedreschi, S. Sethi, B. Harris, S. Greenstreet, N. Wolf, S. Smeltz, and C. McGonigle. 2020. Combining fisheries surveys to inform marine species distribution modelling. ICES Journal of Marine Science 77(2): 539–552.

Moyle, P.B., and D.K. Stompe. 2022. Non-native fishes in estuaries. In Fish and fishes in estuaries: A global perspective, ed A.K. Whitfield, K.W. Able, S.J.M. Blaber, and M. Elliott, 684–705. Hoboken: Wiley.

Moyle, P.B., J.A. Hobbs, and J.R. Durand. 2018. Delta Smelt and water politics in California. Fisheries 43(1): 42–50.

Moyle P.B. 2002. Inland fishes of California: Revised and expanded. Berkeley: University of California Press.

Moyle, P.B., B. Herbold, D.E. Stevens, and L.W. Miller. 1992. Life history and status of Delta Smelt in the Sacramento-San Joaquin Estuary. Transactions of the American Fisheries Society 121(1): 67–77.

O’Rear, T.A., J. Montgomery, P.B. Moyle, and J.R. Durand. 2021. Trends in fish and invertebrate populations of Suisun Marsh January 2020 - December 2020. Report. Davis: University of California, Davis.

Pérez-Ruzafa, A., C. Marcos, I.M. Pérez-Ruzafa, and M. Pérez-Marcos. 2011. Coastal lagoons: “transitional ecosystems” between transitional and coastal waters. Journal of Coastal Conservation 15(3): 369–392.

R Core Team. 2020. R: A language and environment for statistical computing. R Foundation for Statistical Computing, Vienna, Austria. https://www.R-project.org/.

Raney, E.C. 1952. The life history of the Striped Bass, Roccus saxatilis (Walbaum). Bulletin of the Bingham Oceanographic Collection 14(1): 5–97.

Sabal, M., S. Hayes, J. Merz, and J. Setka. 2016. Habitat alterations and a nonnative predator, the Striped Bass, increase native Chinook Salmon mortality in the Central Valley, California. North American Journal of Fisheries Management 36(2): 309–320.

Schoellhamer, D.H., S.A. Wright, and J.Z. Drexler. 2013. Adjustment of the San Francisco Estuary and watershed to decreasing sediment supply in the 20th century. Marine Geology 345: 63–71.

Sommer, T., C. Armor, R. Baxter, R. Breuer, L. Brown, M. Chotkowski, S. Culberson, F. Feyrer, M. Gingras, B. Herbold, W. Kimmerer, A. Mueller-Solger, M. Nobriga, and K. Souza. 2007. The collapse of pelagic fishes in the upper San Francisco Estuary. Fisheries 32(6): 270–277.

Stevens, D.E., D.W. Kohlhorst, L.W. Miller, and D.W. Kelley. 1985. The decline of Striped Bass in the Sacramento-San Joaquin Estuary, California. Transactions of the American Fisheries Society 114(1): 12–30.

Stompe, D.K., P.B. Moyle, A. Kruger, and J.R. Durand. 2020. Comparing and integrating fish surveys in the San Francisco Estuary: why diverse long-term monitoring programs are important. San Francisco Estuary and Watershed Science. 18(2).

Thorson, J.T., M.L. Pinsky, and E.J. Ward. 2016. Model-based inference for estimating shifts in species distribution, area occupied and centre of gravity. Methods in Ecology and Evolution 7(8): 990–1002.

Turner, J.L., and H.K. Chadwick. 1972. Distribution and abundance of young-of-the-year Striped Bass, Morone saxatilis, in relation to river flow in the Sacramento-San Joaquin Estuary. Transactions of the American Fisheries Society 101(3): 442–452.

Tweedie, M.C.K. 1984. An index which distinguishes between some important exponential families. In Statistics: Applications and New Directions, Proceedings of the Indian Statistical Institute Golden Jubilee International Conference, 579–604. Calcutta: Indian Statistical Institute.

Vanderstukken, M., N. Mazzeo, W.V. Colen, S.A.J. Declerck, and K. Muylaert. 2011. Biological control of phytoplankton by the subtropical submerged macrophytes Egeria densa and Potamogeton illinoensis: a mesocosm study. Freshwater Biology 56(9): 1837–1849.

Walker, N.D., D.L. Maxwell, W.J.F. Le Quesne, and S. Jennings. 2017. Estimating efficiency of survey and commercial trawl gears from comparisons of catch-ratios. ICES Journal of Marine Science 74(5): 1448–1457.

Whipple, A., R. Grossinger, D. Rankin, B. Stanford, and R. Askevold. 2012. Sacramento-San Joaquin Delta historical ecology investigation: exploring pattern and process. Report 672. Richmond, CA: San Francisco Estuary Institute - Aquatic Science Center.

White, J. 2020. Fall midwater trawl survey end of season report: 2020. Report. Stockton, CA: California Department of Fish and Wildlife.

Whitfield, A.K., K.W. Able, S.J.M. Blaber, M. Elliot, A. Franco, T.D. Harrison, I.C. Potter, and J.R. Tweedley. 2022. Fish assemblages and functional groups. In Fish and fishes in estuaries: A global perspective, ed A.K. Whitfield, K.W. Able, S.J.M. Blaber, and M. Elliott, 16–59. Hoboken: Wiley.

Wickham, H. 2016. ggplot2: Elegant graphics for data analysis. Springer-Verlag New York. Wilson, J.G., 1988. The estuary as a resource. In The Biology of Estuarine Management, 9–27. Dordrecht: Springer.

Wood, S.N. 2011. Fast stable restricted maximum likelihood and marginal likelihood estimation of semiparametric generalized linear models. Journal of the Royal Statistical Society 73(1): 3–36

Yarrow, M., V.H. Marín, M. Finlayson, A. Tironi, L.E. Delgado, and F. Fischer. 2009. The ecology of Egeria densa Planchón (Liliopsida: Alismatales): A wetland ecosystem engineer? Revista chilena de historia natural 82: 299–313.

Yoshiyama, R.M., P.B. Moyle, E.R. Gerstung, and F.W. Fisher. 2000. Chinook Salmon in the California Central Valley: An assessment. Fisheries 25(2): 6–20.

