## Supplemental tables and figures for "A Spatiotemporal History of Key San Francisco Estuary Pelagic Fish Species"

### Supplementary Material:

*Appendix Table 1. Results of DHARMa (Hartig 2022) testing of fixed effect residual uniformity, dispersion, and outliers. Uniformity was tested using One-Sample Kolmogorov-Smirnov test, dispersion using DHARMa nonparametric dispersion test via standard deviation of residuals fitted versus simulated, and outliers using DHARMa outlier test based on exact binomial test with approximate expectations. Disp. = dispersion value and Num = number of outliers detected at both margins. Significant test results at <0.05 level indicated by *. Significant test results do not necessarily indicate misspecification or poor model fit due to the exceptionally large number of data points. See main text for explanation.*

| | Uniformity | | Dispersion | | Outliers | |
|---|---|---|---|---|---|---|
| | **D** | **p-value** | **Disp.** | **p-value** | **Num** | **p-value** |
| ***Striped Bass*** | 0.0041 | 0.0670 | 1.0010 | 0.6600 | 339 | 0.0002* |
| ***Delta Smelt*** | 0.0033 | 0.2010 | 0.9999 | 0.9880 | 385 | 0.1828 |
| ***Longfin Smelt*** | 0.0033 | 0.2108 | 0.9998 | 0.9920 | 386 | 0.1995 |
| ***Threadfin Shad*** | 0.0024 | 0.5877 | 0.9964 | 0.5360 | 371 | 0.0406* |
| ***American Shad*** | 0.0038 | 0.1087 | 0.9915 | 0.0120* | 467 | 0.0083* |

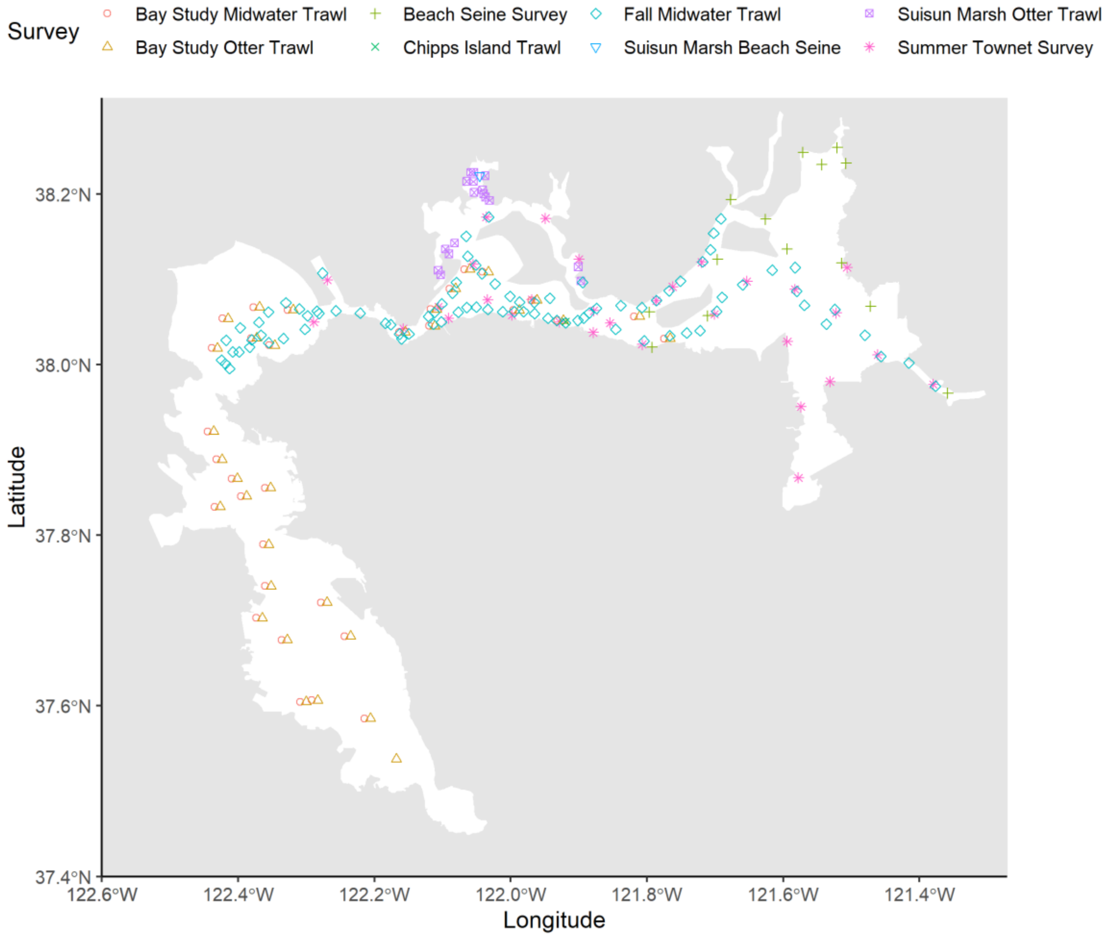


*Appendix Figure 1. Location of sampling stations included in generalized linear mixed models of species occurrence. Different colors/shapes represent the eight surveys.*

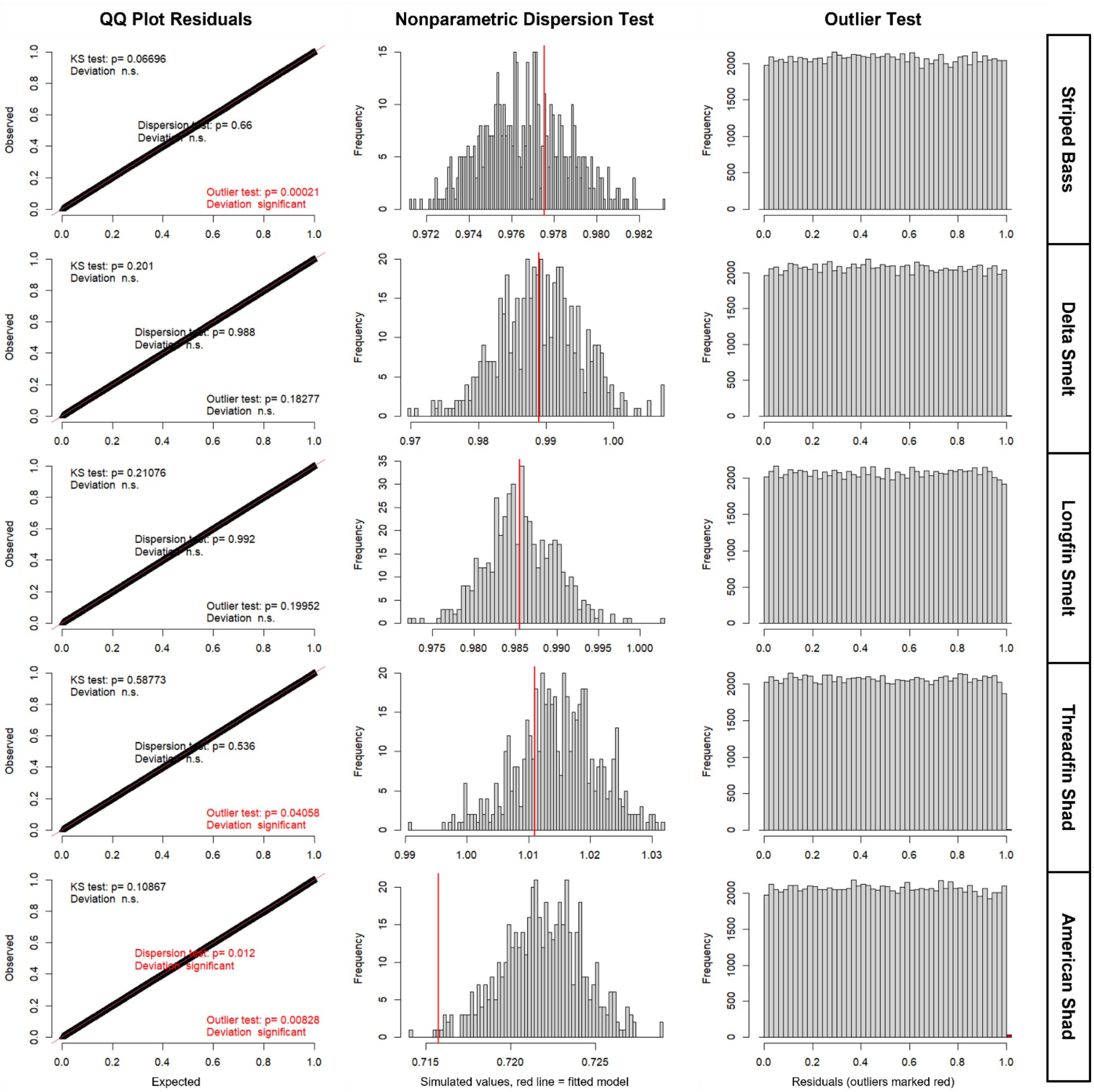


*Appendix Figure 2. Model diagnostics from DHARMa of fixed effect residuals from binomial GLMMs. Left column is QQ plots of observed versus expected values, middle column is standard deviation of fitted versus simulated residuals, and right column is residual values with outliers marked in red. Uniformity was tested using One-Sample Kolmogorov-Smirnov test, dispersion using DHARMa nonparametric dispersion test via standard deviation of residuals fitted versus simulated, and outliers using DHARMa outlier test based on exact binomial test with approximate expectations.*
